# Mitochondrial fusion and altered beta-oxidation drive muscle wasting in a *Drosophila* cachexia model

**DOI:** 10.1101/2023.06.23.546217

**Authors:** Callum Dark, Nashia Ali, Sofia Golenkina, Ronnie Blazev, Benjamin L Parker, Katherine Murphy, Gordon Lynch, Tarosi Senapati, S Sean Millard, Sarah M Judge, Andrew R Judge, Louise Y Cheng

## Abstract

Cancer cachexia is a tumour-induced wasting syndrome, characterised by extreme loss of skeletal muscle. Defective mitochondria can contribute to muscle wasting; however, the underlying mechanisms remain unclear. Using a *Drosophila* larval model of cancer cachexia, we observed enlarged and dysfunctional muscle mitochondria. Morphological changes were accompanied by upregulation of beta-oxidation proteins and depletion of muscle glycogen and lipid stores. Muscle lipid stores were also decreased in Colon-26 adenocarcinoma mouse muscle samples, and expression of the beta-oxidation gene *CPT1A* was negatively associated with muscle quality in cachectic patients. Mechanistically, mitochondrial defects result from reduced muscle insulin signalling, downstream of tumour-secreted insulin growth factor binding protein (IGFBP) homolog ImpL2. Strikingly, muscle-specific inhibition of Forkhead box O (FOXO), mitochondrial fusion, or beta-oxidation in tumour-bearing animals preserved muscle integrity. Finally, dietary supplementation with nicotinamide or lipids, improved muscle health in tumour-bearing animals. Overall, our work demonstrates that muscle FOXO, mitochondria dynamics/beta-oxidation and lipid utilisation are key regulators of muscle wasting in cancer cachexia.

## Introduction

Cancer cachexia is a metabolic wasting syndrome characterised by the involuntary loss of muscle and adipose tissue, caused by tumour-secreted factors, and is the cause of up to 20% of cancer-related deaths^1^. The most prominent clinical feature of cachexia is the continuous loss of skeletal muscle, which cannot be fully reversed by conventional nutritional support.

*Drosophila melanogaster* is emerging as an excellent model to identify tumour-secreted factors that drive cancer cachexia. Adult and larval tumour models that induce cachectic phenotypes have been established in *Drosophila*, and these models have revealed several mechanisms by which tumours induce these phenotypes^2–11^. In mouse and *Drosophila* models of cachexia, it has been shown that tumours can induce a number of metabolic alterations in the muscle^12,13^, such as increased autophagy^4^, proteolysis^5^, decreased protein synthesis^4,5^, defective mitochondrial function^2,8^, reduced ATP production^2,8^ and depleted Extra Cellular Matrix (ECM)^4^. While all these changes are symptomatic of cachexia, it is not clear whether they directly drive cachexia; furthermore, it is unclear how these defects are linked to tumour-secreted factors.

In this study, utilising our previously characterised eye imaginal disc tumour models^4^, we identified two muscle-specific mechanisms that contribute to muscle wasting: 1) altered activity of the transcription factor Forkhead box O (FOXO), a negative regulator of insulin signalling and 2) increased beta-oxidation resulting from mitochondria fusion. Both mechanisms are mediated by a reduction of systemic insulin signalling induced by tumour-secreted insulin-like-peptide antagonist ImpL2 (Imaginal morphogenesis protein-Late 2). Downstream of FOXO, mitochondrial fusion (mediated by Mitochondrial assembly regulatory factor (Marf)) and increased beta-oxidation (mediated by Withered (Whd)) were responsible for increased utilisation of muscle lipids. Strikingly, inhibiting FOXO, Marf or Whd specifically in the muscle was sufficient to over-ride the effects of tumour-secreted factors and improve muscle morphology when tumours were present. In addition, feeding cachectic animals a diet supplemented with nicotinamide (Vitamin B3), or a high-fat coconut oil diet, which replenished lipids in the muscle, were sufficient to improve muscle integrity. Finally, we show that these processes are important in other cachexia models. We observed a similar depletion of lipid reserves in a C-26 mouse cachexia model; furthermore, beta-oxidation gene Carnitine Palmitoyltransferase 1A (*CPT1A*, the mammalian homolog of Whd) is negatively correlated with muscle quality in cachectic patients with pancreatic ductal adenocarcinoma. Together, our study shows that mitochondrial fusion, beta-oxidation and lipid utilisation are likely early hallmarks of muscle disruption during cancer cachexia.

## Results

### Inhibition of mitochondrial fusion prevents muscle detachment in tumour-bearing animals

Studies examining the muscles of cachectic patients have demonstrated increased mitochondrial energy expenditure^14^ and size^14,15^, however, how changes in mitochondrial morphology underly cachexia is so far not clear. In this study, we utilise two *Drosophila* larval tumour models to study muscle biology. In the first model, the tumour is induced via the GAL4-UAS mediated overexpression of *Ras^V^*^12^ and Disc Large (Dlg) RNAi in the eye (Figure 1 A). In the second model, the tumour is induced via the QF2-QUAS mediated overexpression of *Ras^V^*^12^ and *scrib* RNAi (Figure 1 A), allowing us to knockdown or overexpress genes of interest in the muscles of tumour-bearing animals using drivers such as *MHC-GAL4* or *mef2-GAL4* (Figure 1 A). Using these models, we have previously shown that tumours caused a loss of muscle integrity^4^.

**Figure 1.**
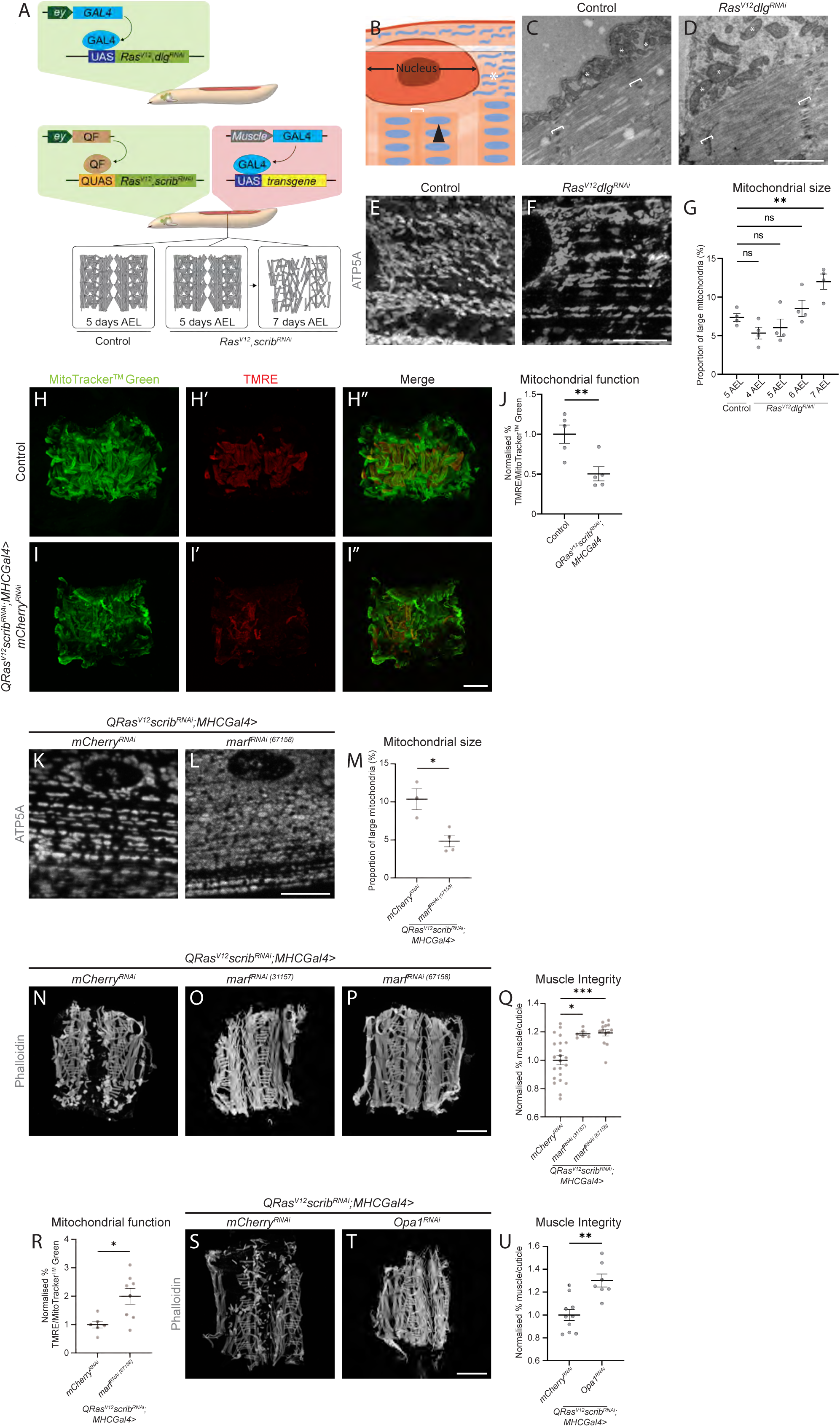
Blocking mitochondrial fusion in cachectic muscles is sufficient to restore muscle mitochondrial defects, as well as muscle health. A) Cartoon schematic depicting the two tumour models used in this study. The first model utilises the GAL4-UAS system to overexpress *Ras^V^*^12^ and *dlg^RNAi^* in the eye discs of *Drosophila* larvae. The second model uses the QF2-QUAS system to overexpress *Ras^V^*^12^ and *scrib^RNAi^* in the eye discs, which allows us to utilise the GAL4-UAS system to simultaneously drive genetic manipulations in the muscle. Both of these systems have comparable muscle integrity to controls at 5 days AEL, but show muscle detachment by 7 days AEL. B) Cartoon representation of a cross section through a larval muscle. The black arrow points to a mitochondrion found in the sarcomere, while white asterisks point to the mitochondria found in the sub-sarcolemma. The white bracket indicates the z-disc of the sarcomere. The transparent white rectangle indicates the plane at which the confocal images and mitochondrial quantifications were acquired in (at the level of the sub-sarcolemma). C, D) Electron microscopy images of mitochondria in the muscles of control and *Ras^V12^dlg^RNAi^* larvae. White asterisks mark mitochondria in the sub-sarcolemma, white brackets indicate z-discs of the sarcomeres. E, F) Representative images of ATP5A staining of mitochondria in the muscles of control and *Ras^V12^dlg^RNAi^*larvae. G) Quantification of the proportion of large mitochondria out of total mitochondria in control and *Ras^V12^dlg^RNAi^* larvae at days 4-7 AEL, performed using One-Way ANOVA (n = 4, 4, 4, 4). H-I’’) Representative images of control and *QRas^V12^scrib^RNAi^;MHC-Gal4>mCherry^RNAi^* larval muscle fillets stained with MitoTracker^TM^ Green which labels all mitochondria (H, I), and active mitochondria with TMRE (H’, I’). Merged images are shown in H’’ and I’’. J) Quantification of the percentage of total mitochondria stained with MitoTracker^TM^ Green that are also positive for TMRE in H’’ and I’’, performed using Student’s t-test (n = 5, 5). K, L) Representative images of ATP5A staining of mitochondria in the muscles of *QRas^V12^scrib^RNAi^;MHC-Gal4>mCherry^RNAi^* and *QRas^V12^scrib^RNAi^;MHC-Gal4>marf^RNAi^* larvae. M) Quantification of the proportion of large mitochondria out of total mitochondria in K and L, performed using Student’s t-test (n = 3, 4). N, O, P) Representative muscle fillets from *QRas^V12^scrib^RNAi^;MHC-Gal4>mCherry^RNAi^*(N), *QRas^V12^scrib^RNAi^;MHC-Gal4>marf^RNAi^* (31157) (O), and *QRas^V12^scrib^RNAi^;MHC-Gal4>marf^RNAi^*(67158) (P) larvae, stained with Phalloidin to visualise actin. Q) Quantification of muscle integrity in N, O and P, performed using Brown-Forsythe (n = 22, 6, 13). R) Quantification of the percentage of total mitochondria stained with MitoTracker^TM^ Green that are shown to be active via TMRE incorporation in the muscles of 6 days AEL *QRas^V12^scrib^RNAi^;MHC-Gal4>mCherry^RNAi^* and *QRas^V12^scrib^RNAi^;MHC-Gal4>marf^RNAi^* (67158) larvae, performed using Student’s t-test (n = 6, 8). S, T) Representative muscle fillets from *QRas^V12^scrib^RNAi^;MHC-Gal4>mCherry^RNAi^* and *QRas^V12^scrib^RNAi^;MHC-Gal4>Opa1^RNAi^*larvae, stained with Phalloidin to visualise actin. U) Quantification of muscle integrity in S and T, performed using Student’s t-test (n = 10, 7). Images were taken at 5 days AEL for (C and E), 6 days AEL for (H, H’, H’’, I, I’, I’’), and 7 days AEL for (D, F, K, L, N, O, P, S and T). Scale bars: 1 μm for (C and D), 10 μm for (E, F, K and L), and 500 μm for (H, H’, H’’, I, I’, I’’, N, O, P, S and T). All error bars are +/- SEM. P values are: ns (not significant), p > 0.05; *, p < 0.05; **, p < 0.01; ***, p < 0.001; ****, p < 0.0001.

Using Electron Microscopy (EM), we observed fewer and larger mitochondria in the sub-sarcolemma of muscles of *Ras^V12^dlg^RNAi^* tumour bearing animals at 7 days after egg lay (AEL, Figure 1 C-D). A disruption of muscle mitochondria morphology in the sub-sarcolemma plane of the muscle (as depicted in a cartoon of a muscle section, Figure 1 B) was further confirmed by antibody staining against mitochondria protein ATP5A, where a significant increase in size was first detected at 7 days AEL (*Ras^V12^dlg^RNAi^*, Figure 1 E-G. For details on mitochondria size quantification see Figure S1). To test whether this increase in mitochondrial size could lead to compromised mitochondrial function, we performed live staining with tetramethylrhodamine ethyl ester (TMRE), a compound used to measure the membrane potential of mitochondria^16^. We detected a significant reduction in TMRE fluorescence in the muscles of tumour-bearing animals (*QRas^V12^scrib^RNAi^*), indicative of reduced membrane potential, and compromised mitochondrial function (Figure 1 H-J). Enlarged mitochondria has been shown to be attributed to mitochondrial fusion, a process evoked to buffer against mitochondrial damage caused by increased reactive oxygen species^17,18^. We found that there was a significant increase in ROS levels (as indicated by dihydroethidium (DHE) staining) in the muscles of tumour-bearing animals (*QRas^V12^scrib^RNAi^*, Figure S2 A-C). To determine if reducing ROS could preserve muscle morphology, we overexpressed ROS scavengers Catalase (Cat) and superoxide dismutase (Sod1), or antioxidative enzyme glutathione peroxidase 1 (GPx1, previously validated in ^19–21)^, specifically in the muscles of tumour-bearing animals (*QRas^V12^scrib^RNAi^*). However, these manipulations did not rescue muscle integrity (Figure S2 D-H, J), suggesting that ROS was a consequence but not the cause of muscle wasting.

Next, we asked if preventing mitochondrial fusion through muscle-specific knockdown of Marf, a protein important for outer mitochondrial membrane fusion^22^, could improve muscle integrity in tumour-bearing animals (*QRas^V12^scrib^RNAi^*). Indeed, Marf knockdown, which effectively reduced mitochondria size (Figure 1 K-M), also significantly improved muscle integrity (Figure 1 N-Q). This manipulation was also able to significantly improve mitochondrial function as assessed by TMRE assay (Figure 1 R). In addition, knockdown of Optic atrophy 1 (Opa1), a protein involved with inner mitochondrial membrane fusion, was also able to improve muscle integrity in tumour-bearing animals (*QRas^V12^scrib^RNAi^,* Figure 1 S-U)^23^. However, overexpression of mitochondrial fission protein Dynamin related protein 1 (Drp1)^23^, did not help preserve muscle integrity (*QRas^V12^scrib^RNAi^*, Figure S2 G, I, J). Together, these experiments suggest that preventing mitochondrial fusion (rather than increasing fission) helps to improve muscle function.

### Reduced insulin signalling mediates changes in mitochondrial size

Mitochondria are known to change in size in response to environmental stressors, especially starvation^24,25^. Upon subjecting wildtype larvae to 24 hrs of nutrient restriction, which is known to reduce systemic insulin signalling^26^, we observed an increase in mitochondrial size (Figure 2 A-C), that is reminiscent of what we saw in the muscles of tumour-bearing animals. We previously reported that *Ras^V12^dlg^RNAi^* eye imaginal disc tumours express elevated levels of ImpL2, and its knockdown ameliorated muscle disruption without affecting tumour size^4^. Here, we found that tumour-specific inhibition of ImpL2 significantly reduced muscle mitochondrial size (*Ras^V12^dlg^RNAi^*, Figure 2 D-F). Together, these data suggest that mitochondrial fusion may be caused by reduced systemic insulin signalling.

**Figure 2.**
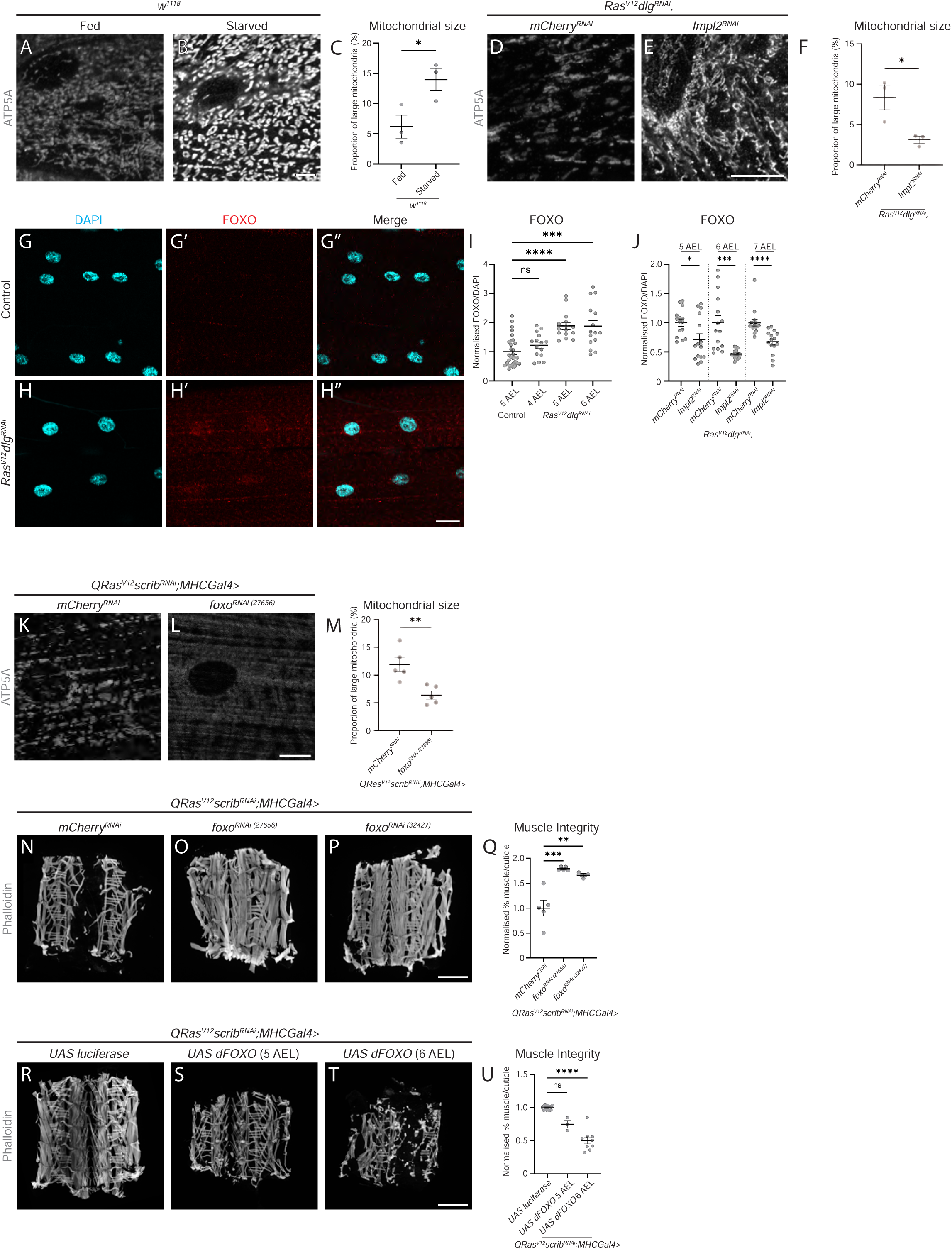
Tumour-secreted Impl2 mediates muscle mitochondrial morphology via FOXO. A, B) Representative images of ATP5A staining of mitochondria in the muscles of *w*^1118^ larvae raised on a normal diet and *w*^1118^ larvae undergoing nutrient restriction from critical weight (60 hr after larval hatching (ALH)). C) Quantification of the proportion of large mitochondria out of total mitochondria in A and B, performed using Student’s t-test (n = 3, 3) D, E) Representative images of ATP5A staining of mitochondria in the muscles of *Ras^V12^dlg^RNAi^,mCherry^RNAi^* and *Ras^V12^dlg^RNAi^,ImpL2^RNAi^* larvae. F) Quantification of the proportion of large mitochondria out of total mitochondria in D and E, performed using Student’s t-test (n = 3, 3). G-H’’) Representative images of control and *Ras^V12^dlg^RNAi^* larval muscles stained with DAPI to label the nucleus (G, H), and FOXO (G’, H’). Merged images are shown in G’’ and H’’. I) Quantification of nuclear FOXO staining in control and *Ras^V12^dlg^RNAi^* larvae at days 5 and 6 AEL performed using Kruskal-Wallis (n = 30, 15, 15, 15). J) Quantification of nuclear FOXO staining of *Ras^V12^dlg^RNAi^,mCherry^RNAi^*and *Ras^V12^dlg^RNAi^,ImpL2^RNAi^* larval muscles at 5, 6 and 7 days AEL, performed using Mann-Whitney U (5 and 7 days AEL), and Welch’s t-test (6 days AEL, n = 15, 15, 15, 15, 15, 15). K, L) Representative images of ATP5A staining of mitochondria in the muscles of *QRas^V12^scrib^RNAi^;MHC-Gal4>mCherry^RNAi^* and *QRas^V12^scrib^RNAi^;MHC-Gal4>foxo^RNAi^* (27656) larvae. M) Quantification of the proportion of large mitochondria out of total mitochondria in K and L, performed using Student’s t-test (n = 5, 5). N, O, P) Representative muscle fillets from *QRas^V12^scrib^RNAi^;MHC-Gal4>mCherry^RNAi^*(N), *QRas^V12^scrib^RNAi^;MHC-Gal4>foxo^RNAi^* (27656) (O), and *QRas^V12^scrib^RNAi^;MHC-Gal4>foxo^RNAi^*(32427) (P) larvae, stained with Phalloidin to visualise actin. Q) Quantification of muscle integrity in N, O and P, performed using One-Way ANOVA (n = 5, 5, 3). R, S, T) Representative muscle fillets from *QRas^V12^scrib^RNAi^;MHC-Gal4>UAS luciferase* (R), *QRas^V12^scrib^RNAi^;MHC-Gal4>UAS-dFOXO* (5 days AEL, S), and *QRas^V12^scrib^RNAi^;MHC-Gal4>UAS-dFOXO* (6 days AEL, T) larvae, stained with Phalloidin to visualise actin. U) Quantification of muscle integrity in R, S and T, performed using Brown-Forsythe (n = 14, 3, 9). Images were taken at 5 days AEL for (A, B, G, G’, G’’ and S), 6 days AEL for (D, E, K, L, R and T), and 7 days AEL for (H, H’, H’’, N, O and P). Scale bars: 10 μm for (A, B, D, E, K and L), 20 μm for (G, G’, G’’, H, H’ and H’’), and 500 μm for (N, O, P, R, S and T). All error bars are +/- SEM. P values are: ns (not significant), p > 0.05; *, p < 0.05; **, p < 0.01; ***, p < 0.001; ****, p < 0.0001.

FOXO transcription factors have been implicated in muscle atrophy in wildtype animals via ubiquitin-proteasome mediated mechanisms^27^ and have been shown to be activated in response to decreased insulin/IGF signalling. In cachectic muscles (*Ras^V12^dlg^RNAi^*), we found a significant increase in nuclear FOXO signal, which first occurred at 5 days AEL (Figure 2 G-I). This preceded muscle disruption at 7 days AEL^4^, suggesting that FOXO may be a key upstream mediator of muscle wasting. Furthermore, tumour-specific ImpL2 inhibition was able to significantly reduce muscle FOXO levels at 5 days AEL (*Ras^V12^dlg^RNAi^*, Figure 2 J).

As the overexpression of FOXO has been linked with increased expression of mitochondrial fusion proteins in mice^28–30^, we next assessed whether increased mitochondrial size in cachectic muscles was regulated by FOXO. Muscle-specific FOXO knockdown was able to reduce mitochondrial size in tumour-bearing animals (*QRas^V12^scrib^RNAi^*, Figure 2 K-M), as well as significantly improve muscle integrity (Figure 2 N-Q). Conversely, overexpression of dFOXO in the muscles of tumour bearing animals (*QRas^V12^scrib^RNAi^*), caused precocious muscle detachment at 6 days AEL (Figure 2 R-U). Together, these results indicate that FOXO plays an important role in regulating mitochondrial morphology as well as muscle integrity downstream of tumour-induced signals.

### Disrupted autophagy and translation in cachectic muscles is not necessary for muscle detachment

Mitochondria disruption has been reported to be accompanied by increased autophagy^31^ and reduced protein translation^32^. Using a transgenic line that expresses a tandem autophagy reporter (UAS pGFP-mCherry-Atg8a)^33^ under the control of a muscle specific driver (*MHC-GAL4*), we found the ratio of mCherry vs. EGFP was significantly elevated in tumour-bearing animals (*QRas^V12^scrib^RNAi^*) at 6 days AEL (Figure S3 A-C). This indicates there is an increased lysosomal degradation and autophagy in the muscles of tumour-bearing animals^34^ prior to the increase in mitochondrial size at day 7. Next, we examined the expression of the sarcomere structural protein Myosin Heavy Chain (MHC), a previously reported proxy for protein synthesis^35^. We found MHC levels were significantly downregulated at 6 days AEL in tumour bearing animals (*Ras^V12^dlg^RNAi^*, Figure S3 D-F). Next, we inhibited tumour-secreted ImpL2 (*Ras^V12^dlg^RNAi^*), and we found that translation levels were significantly restored by 5 days AEL (Figure S3 J-L), and autophagy levels were restored by 7 days AEL (Figure S3 G-I), confirming that muscle perturbations are driven by tumour-secreted ImpL2.

As disruptions to autophagy and protein synthesis preceded muscle detachment (at 7 days AEL^4^), we functionally assessed whether inhibiting autophagy or enhancing translation could rescue muscle integrity (*QRas^V12^scrib^RNAi^*). However, neither the expression of a constitutively activated S6 kinase (S6K^CA^), which has previously been shown to sufficiently increase protein translation in the muscle^35^, nor the inhibition of autophagy via the expression of a RNAi against the protein kinase Atg1 which has previously been shown to reduce autophagy^36^, was able to prevent tumour-induced muscle detachment (Figure S3 M-R). Due to the role of autophagy as a stress compensation mechanism^37^, we also tried increasing autophagy specifically in the muscles through the overexpression of Atg1. However, this led to lethality of the larvae at the 1^st^ instar (data not shown). Together this data suggests that defects in autophagy and protein synthesis are not the primary cause of muscle wasting.

### Mitochondrial fusion promotes lipid utilisation in cachectic muscles

To further investigate the role of insulin-mediated mitochondrial fusion on muscle wasting, we next examined how these changes affected lipid and glycogen metabolism in the muscle. Cachexia has been associated with the rewiring of lipid metabolism^38^, and we have previously shown that there is a significant reduction in total lipid stores of cachectic animals^4^. Consistent with a reduction in overall lipid stores, we observed a significant reduction in the number of lipid droplets (LDs, Figure 3 A-C and Figure S4 A-B, visualised by LipidTOX^TM^), in the muscles of tumour bearing animals (*Ras^V12^dlg^RNAi^*). The depletion of lipids is phenocopied by nutrient restriction in wildtype animals (Figure 3 D-F). Furthermore, inhibition of ImpL2 in the tumour was sufficient to significantly increase muscle LD levels by 6 days AEL (*Ras^V12^dlg^RNAi^*, Figure 3 G-I). Together, our data suggests that the depletion of muscle lipid stores lies downstream of insulin signalling.

**Figure 3.**
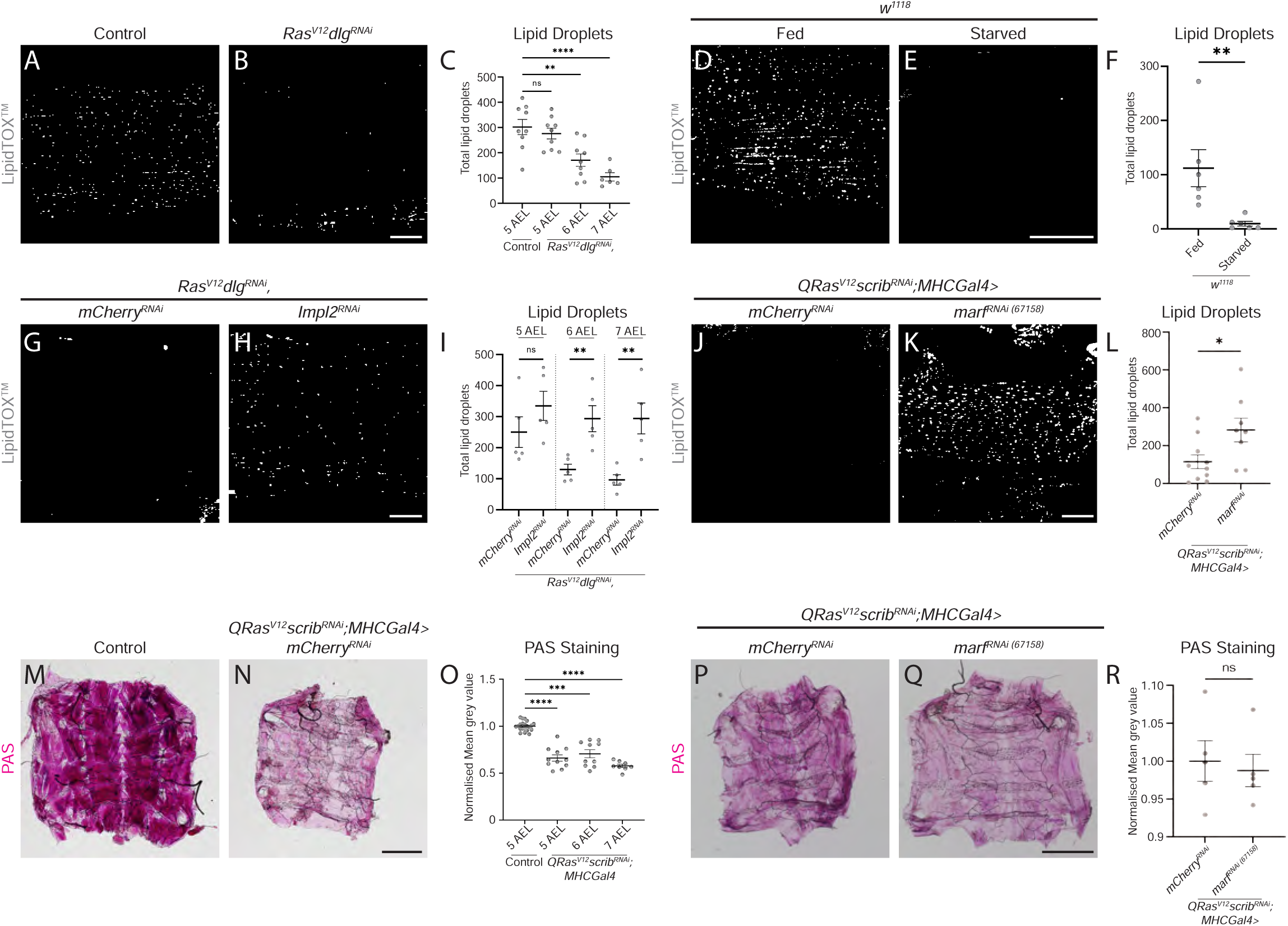
Blocking mitochondrial fusion restores muscle lipid stores, but not glycogen stores. A, B) Binary representation of lipid droplets (LDs) stained with LipidTOX^TM^ in the muscles of control and *Ras^V12^dlg^RNAi^* larvae from S4 A, B, obtained through thresholding in FIJI. More details on thresholding can be found in the Methods and Materials. C) Quantification of the number of LDs in control and *Ras^V12^dlg^RNAi^* larvae at days 5-7 AEL, performed using One-Way ANOVA (n = 9, 9, 9, 6). D, E) Binary images of LDs stained with LipidTOX^TM^ in the muscles of *w*^1118^ larvae raised on a normal diet and *w*^1118^ larvae undergoing nutrient restriction from critical weight (60 hr after larval hatching (ALH)), obtained through thresholding in FIJI. F) Quantification of the number of LDs in D and E, performed using Mann-Whitney U (n = 6, 6). G, H) Binary representation of LDs stained with LipidTOX^TM^ in the muscles of *Ras^V12^dlg^RNAi^,mCherry^RNAi^* and *Ras^V12^dlg^RNAi^,ImpL2^RNAi^* larvae obtained through thresholding in FIJI. I) Quantification of the number of LDs in G and H, as well as from earlier timepoints, performed using Student’s t-tests (n = 5, 5, 5, 5, 5, 5). J, K) Binary representation of LDs stained with LipidTOX^TM^ in the muscles of *QRas^V12^scrib^RNAi^;MHC-Gal4>mCherry^RNAi^* and *QRas^V12^scrib^RNAi^;MHC-Gal4>marf^RNAi^* (67158) larvae, obtained through thresholding in FIJI. L) Quantification of the number of LDs in J and K, performed using Student’s t-test (n = 10, 8). M, N) Representative images of muscle fillets from control and *QRas^V12^scrib^RNAi^;MHC-Gal4>mCherry^RNAi^* larvae, stained with periodic acid solution (PAS) to visualise polysaccharides such as glycogen. O) Quantification of PAS staining in control and *QRas^V12^scrib^RNAi^;MHC-Gal4>mCherry^RNAi^* at days 5-7 AEL using Brown-Forsythe (n = 21, 11, 10, 8). P, Q) Representative images of muscle fillets from *QRas^V12^scrib^RNAi^;MHC-Gal4>mCherry^RNAi^* and *QRas^V12^scrib^RNAi^;MHC-Gal4>marf^RNAi^* (67158) larvae, stained with periodic acid solution (PAS) to visualise polysaccharides such as glycogen. R) Quantification of PAS staining in P and Q, performed using Student’s t-test (n = 5, 5). Images were taken at 5 days AEL for (A, D, E and M), and 7 days AEL for (B, G, H, J, K, N, P and Q). Scale bars: 20 μm for (A, B, D, E, G, H, J and K), and 500 μm for (M, N, P and Q). All error bars are +/- SEM. P values are: ns (not significant), p > 0.05; *, p < 0.05; **, p < 0.01; ***, p < 0.001; ****, p < 0.0001.

The lack of LDs in the muscle could be accounted for by an increase in lipolysis, whereby fat storage is broken down and released as free fatty acids. However, we found that by promoting LD formation through the overexpression of lipid storage droplet-2 (Lsd-2, Figure S4 C-E) did not prevent the loss of muscle integrity in tumour-bearing animals (*QRas^V12^scrib^RNAi^*, Figure S4 F-H).

Next, we tested if inhibiting mitochondria fusion via Marf RNAi could restore lipid stores. We found that Marf knockdown in the muscles of tumour-bearing animals restored LD numbers (*QRas^V12^scrib^RNAi^*, Figure 3 J-L). It is known that utilisation of glycogen, which is a readily available source of energy, often occurs before fat breakdown^39^. Consistent with this, we found that glycogen stores in the muscles were depleted earlier than lipid stores at 5 days AEL (*Ras^V12^dlg^RNAi^*, Figure 3 M-O). However, muscle-specific knockdown of Marf was not able to preserve the loss of muscle glycogen stores (*QRas^V12^scrib^RNAi^*, Figure 3 P-R), suggesting that mitochondrial fusion had greater effects on lipid rather than glycogen stores in the muscles of cachectic animals.

### Inhibition of beta-oxidation improves muscle integrity

To better understand the metabolic changes that occur in the muscles of tumour-bearing animals, we conducted proteomics to assess for protein expression levels in wildtype and tumour-bearing (*Ras^V12^dlg^RNAi^*) muscles. Consistent with our immunostaining results (Figure S3 D-F), where we observed a significant downregulation of MHC levels, we detected a global downregulation of translational proteins (Figure 4 A). Furthermore, we found that proteins involved in the GO term “metabolism of lipids” were significantly upregulated (Figure 4 A-B). Interestingly, CPT1A/Whd, an important regulator of the beta-oxidation pathway^40^, was found to be upregulated both at the transcriptional (*QRas^V12^scrib^RNAi^*, Figure 4 C) and protein levels (*Ras^V12^dlg^RNAi^*, Figure 4 D) in the muscles of tumour-bearing animals. Beta-oxidation is the metabolic process by which fatty acids are broken down into acetyl-CoA molecules, which can then be used to produce energy (Figure 4 E). As beta-oxidation proceeds, the fatty acids stored in the LDs become depleted, leading to a reduction in the size and abundance of LDs in the cell. To test if preventing beta-oxidation could affect muscle integrity, we next knocked down Whd specifically in the cachectic muscles (*QRas^V12^scrib^RNAi^*). We found that Whd inhibition was sufficient to significantly increase LD numbers (Figure 4 F) and improve muscle integrity (Figure 4 G-I) by 7 days AEL, suggesting that beta-oxidation is a key regulator of muscle integrity in cachectic muscles. Interestingly, while Whd knockdown was able to restore mitochondrial function (as assessed via TMRE, Figure 4 J), it was however not able to restore mitochondrial size (Figure 4 K-M). To assess if *whd* is regulated by insulin signalling, we knocked down FOXO via RNAi, and found that this was able to inhibit the transcription of *whd* (*QRas^V12^scrib^RNAi^*, Figure 4 C). Together, this data suggests that FOXO lies upstream of beta-oxidation, and mitochondria function lies downstream of beta-oxidation.

**Figure 4.**
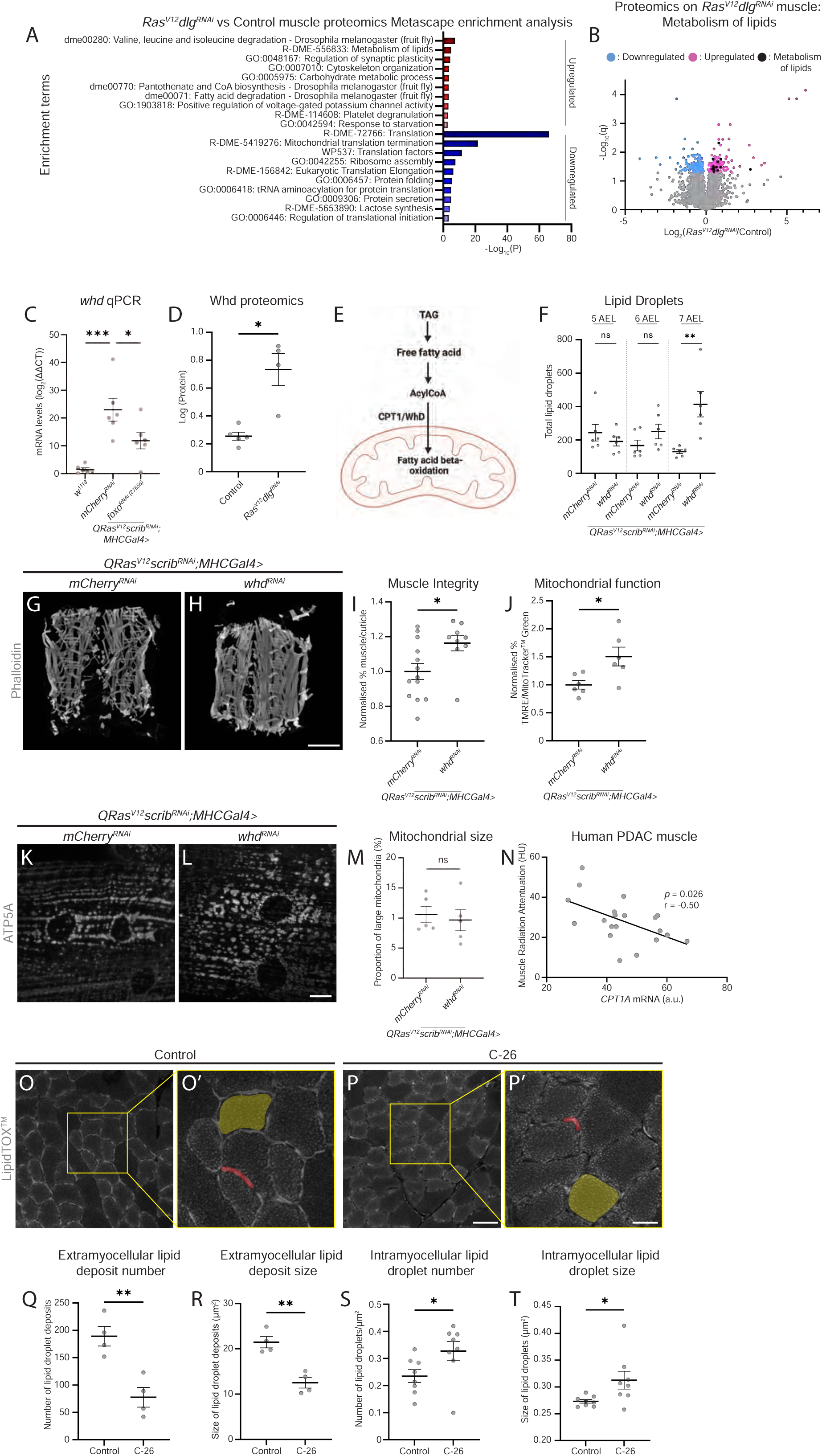
Lipid metabolism via beta-oxidation is a mediator of muscle wasting. A) Proteomics were performed using the muscles from 5 days AEL control and 7 days AEL *Ras^V12^dlg^RNAi^*larvae (n = 5, 4). The top ten up and downregulated enrichment terms from Metascape enrichment analysis are displayed. B) Volcano plot showing differentially regulated proteins in control vs *Ras^V12^dlg^RNAi^* larval muscles, with proteins found in the “R-DME-556833: Metabolism of lipids” enrichment term highlighted (n= 5, 4). C) Quantification of qPCR results examining whd mRNA levels in *w*^1118^, *QRas^V12^scrib^RNAi^;MHCGal4>mCherry^RNAi^*, and *QRas^V12^scrib^RNAi^;MHCGal4>foxo^RNAi^* (27656) larval muscles at 6 days AEL, performed using One-Way ANOVA (n = 6, 6, 6) D) Quantification of Log transformed whd protein levels, from proteomics in B, performed using Mann-Whitney U (n = 5, 4). E) Schematic detailing Whd’s role in mitochondrial beta-oxidation. F) Quantification of the number of lipid droplets in *QRas^V12^scrib^RNAi^;MHC-Gal4>mCherry^RNAi^* and *QRas^V12^scrib^RNAi^;MHC-Gal4>whd^RNAi^* larval muscles, at 5, 6 and 7 days AEL, performed using Mann-Whitney U (5 and 6 days AEL) and Student’s t-test (7 days AEL, n = 6, 6, 6, 6). G, H) Representative muscle fillets from *QRas^V12^scrib^RNAi^;MHC-Gal4>mCherry^RNAi^*and *QRas^V12^scrib^RNAi^;MHC-Gal4>whd^RNAi^* larvae, stained with Phalloidin to visualise actin. I) Quantification of muscle integrity in G and H, performed using Student’s t-test (n = 13, 9). J) Quantification of the percentage of total mitochondria stained with MitoTracker^TM^ Green that are shown to be active via TMRE incorporation in the muscles of 6 days AEL *QRas^V12^scrib^RNAi^;MHC-Gal4>mCherry^RNAi^* and *QRas^V12^scrib^RNAi^;MHC-Gal4>whd^RNAi^* larvae, performed using Student’s t-test (n = 6, 6). K, L) Representative images of ATP5A staining of mitochondria in the muscles of *QRas^V12^scrib^RNAi^;MHC-Gal4>mCherry^RNAi^* and *QRas^V12^scrib^RNAi^;MHC-Gal4>whd^RNAi^* larvae. M) Quantification of the proportion of large mitochondria out of total mitochondria in K and L, performed using Student’s t-test (n = 5, 5). N) Quantification of human PDAC muscle CPT1A mRNA levels plotted against Muscle Radiation attenuation, performed using Non-parametric Spearman’s Correlation (n = 20). (a.u.: average unit). O, O’, P, P’) Representative images of cross-sections of Type IIa muscle fibres in mouse tibiallis anterior muscle, stained with LipidTOX^TM^, taken from control (O) and C-26 tumour (P) animals. Yellow boxes highlight zoomed in areas to better visualise intramyocellular and extramyocellular lipid deposits. Examples of extramyocellular lipid droplets have been labelled with red shading. Examples of where intramyocellular lipid droplets were measured have been labelled with yellow shading. Q) Quantification of the number of extramyocellular lipid deposits in O and P, performed using Student’s t-test, (n = 4, 4). R) Quantification of the size of extramyocellular lipid deposits in O and P, performed using Student’s t-test, (n = 4, 4). S) Quantification of the density of intramyocellular lipid deposits in O and P, performed using Student’s t-test, (n = 4, 4). T) Quantification of the size of intramyocellular lipid deposits in O and P, performed using Student’s t-test, (n = 4, 4) Images were taken at 6 days AEL for (K and L), 7 days AEL for (G and H), and 12 weeks plus 17-25 days after subcutaneous injection of PBS or C-26 tumour cells for (O, O’, P and P’). Scale bars: 10 μm for (K and L), 20 μm for (O’ and P’), 50 μm for (O and P), and 500 μm for (G and H). All error bars are +/- SEM. P values are: ns (not significant), p > 0.05; *, p < 0.05; **, p < 0.01; ***, p < 0.001; ****, p < 0.0001.

To assess if there is an association between lipid metabolism and cachexia in patient samples, we examined the relationship between muscle *CPT1A* mRNA levels and muscle radiation attenuation (an indicator of muscle quality), using a microarray data set from patients with pancreatic ductal adenocarcinoma (PDAC)^41^. We found a significant negative correlation of these two parameters (Figure 4 N), suggesting that increased *CPT1A* is correlated with poor muscle health in cachectic PDAC patients.

Finally, we also examined whether lipid metabolism plays a role in the Colon-26 (C-26) xenograft model in the mouse. Cross-sections through muscle samples of control vs. C-26 mice demonstrated a significant depletion of lipid stores, as indicated by a significant decrease in extramyocellular LD number and size (Figure 4 O-R, shaded in red). Interestingly, when we examined intramyocellular LD levels, we observed an increase in density and size of LDs in cachectic muscles compared to control (Figure 4 O-P’, S, T, shaded in yellow), which is consistent with reports of intramyocellular LD accumulation reported in human cachectic muscles^42,43^. Overall, this data suggests that the depletion of muscle lipid stores via beta-oxidation affects mitochondrial function and is negatively correlated with muscle health in cachectic flies, mice and patients.

### Modulation via dietary supplementation of nicotinamide or a high-fat diet improves muscle integrity

To determine whether we could prevent mitochondrial damage and muscle disruption in tumour-bearing animals through dietary modulation, we fed cachectic animals with diets supplemented with nicotinamide or coconut oil. Nicotinamide is a precursor of Vitamin B3 and is known to improve mitochondria health^44^. Tumour-bearing animals (*QRas^V12^scrib^RNAi^*) fed a diet containing 1 g/kg of nicotinamide exhibited reduced muscle mitochondrial size (Figure 5 A-C), FOXO levels (Figure 5 D), and improved muscle integrity (Figure 5 E-G). Coconut oil has previously been shown to enhance overall lipid levels in *Drosophila* larvae^45^. We found this high-fat diet significantly improved LD levels in the muscles of tumour-bearing animals (*Ras^V12^dlg^RNAi^*, Figure 5 H-J). Most excitingly, this dietary supplementation significantly improved muscle integrity (Figure 5 K-M), as well as mitochondrial function (Figure 5 N). Together, this data demonstrates that dietary supplementation of lipids or nicotinamide can potentially improve outcomes in cachexia through modulation of FOXO, lipid metabolism or mitochondrial dynamics.

**Figure 5.**
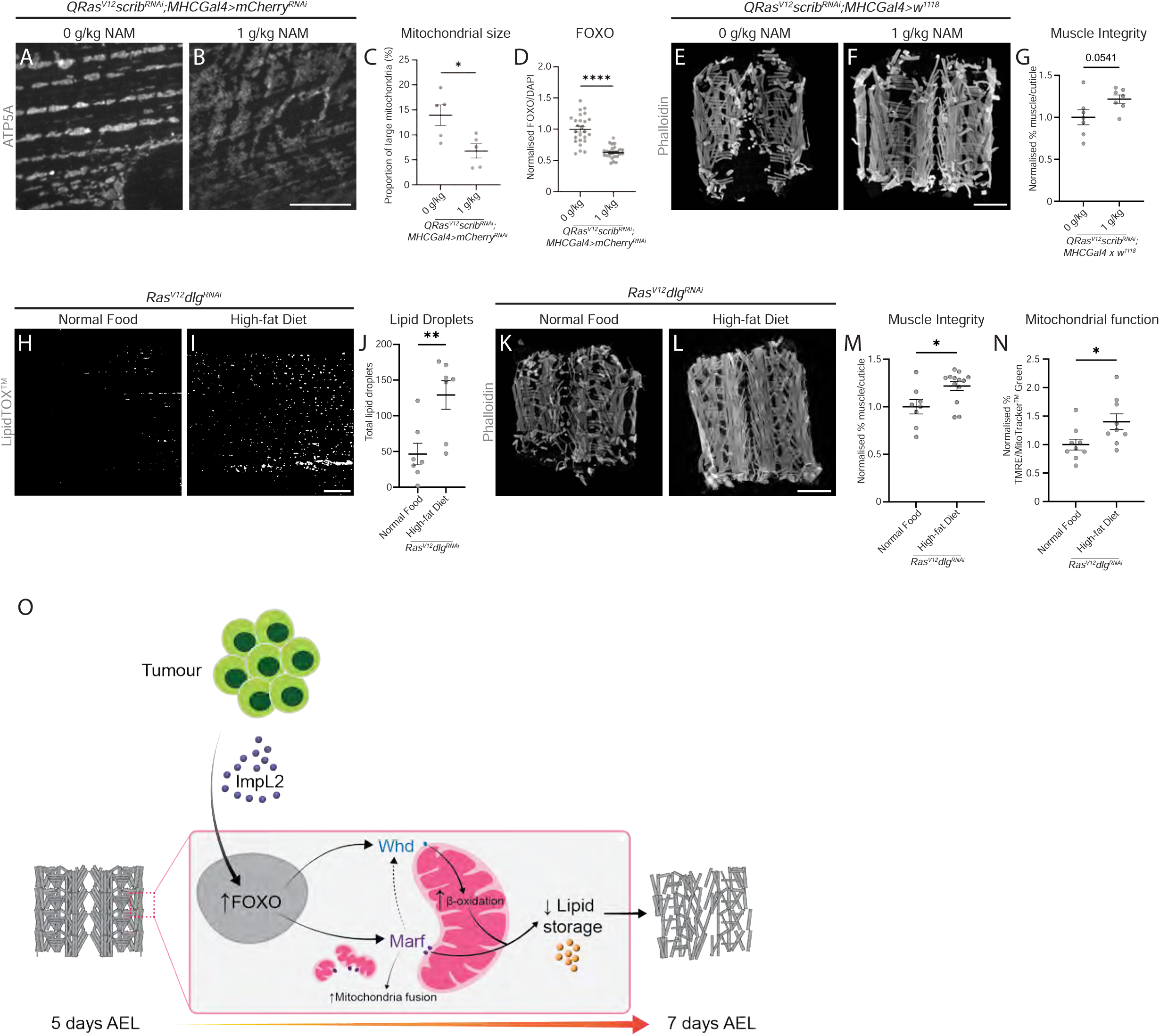
Nicotinamide and high fat diets improve muscle health in tumour-bearing animals. A, B) Representative images of ATP5A staining of mitochondria in the muscles of *QRas^V12^scrib^RNAi^;MHC-Gal4>mCherry^RNAi^* larvae raised on a normal diet, or raised on a diet containing 1 g/kg nicotinamide (NAM). C) Quantification of the proportion of large mitochondria out of total mitochondria in A and B, performed using Student’s t-test (n = 5, 5). D) Quantification of nuclear FOXO staining in the muscles of 6 days AEL *QRas^V12^scrib^RNAi^;MHC-Gal4>mCherry^RNAi^* larvae raised on a normal diet, or raised on a diet containing 1 g/kg nicotinamide (NAM), performed using Welch’s t-test (n = 25, 25). E, F) Representative muscle fillets from *QRas^V12^scrib^RNAi^;MHC-Gal4>w*^1118^ larvae raised on a normal diet, or raised on a diet containing 1 g/kg nicotinamide (NAM), stained with Phalloidin to visualise actin. G) Quantification of muscle integrity in E and F, performed using Student’s t-test (n = 7, 7). H, I) Binary representation of lipid droplets (LDs) stained with LipidTOX^TM^ in the muscles of *Ras^V12^dlg^RNAi^* larvae raised on a normal diet, and raised on a high fat diet, obtained through thresholding in FIJI. J) Quantification of the number of LDs in H and I, performed using Mann-Whitney U (n = 7, 7). K, L) Representative muscle fillets from *Ras^V12^dlg^RNAi^* larvae raised on a normal diet, or raised on a high fat diet, stained with Phalloidin to visualise actin. M) Quantification of muscle integrity in K and L, performed using Mann-Whitney U (n = 8, 13). N) Quantification of the percentage of total mitochondria stained with MitoTracker^TM^ Green that are shown to be active via TMRE incorporation in the muscles of 6 days AEL *Ras^V12^dlg^RNAi^* larvae raised on a normal diet, or raised on a high fat diet, performed using Student’s t-test (n = 9, 9). O) Tumour-secreted ImpL2 mediates insulin signalling in the muscle of tumour-bearing animals by influencing the nuclear localisation of FOXO. This reduction in muscle insulin signalling influences mitochondria fusion via Marf. Increased mitochondrial fusion is accompanied by a decrease in muscle lipid stores, which is the result of increased fatty-acid beta-oxidation via Whd in the mitochondria. This results in muscles that have depleted their energy reserves, contributing to a loss of muscle integrity. Images were taken at 6 days AEL for (H and I), and 7 days AEL for (A, B, D, E, K and L). Scale bars: 10 μm for (D and E), 20 μm for (H and I), and 500 μm for (A, B, K and L). All error bars are +/- SEM. P values are: ns (not significant), p > 0.05; *, p < 0.05; **, p < 0.01; ***, p < 0.001; ****, p < 0.0001.

## Discussion

Utilising a *Drosophila* model of cachexia, we have demonstrated that muscle wasting in cachexia is mediated by two main mechanisms: insulin signalling via FOXO and beta-oxidation via mitochondrial fusion (Figure 5 O). First, tumour-secreted ImpL2 systemically decreases insulin signalling, which results in an increase in FOXO nuclear-localisation in the muscle. This is accompanied by an increase in mitochondrial size and a depletion of the muscle lipid stores. Knockdown of tumour-secreted ImpL2, or muscle-specific FOXO inhibition, resulted in reduced mitochondrial size and improved muscle integrity. Furthermore, inhibition of mitochondrial fusion via Marf, or beta-oxidation via Whd, were also able to improve mitochondrial function and integrity, as well as increase muscle lipid stores. Finally, supplementing dietary lipids through a high-fat diet was sufficient to significantly enhance muscle integrity.

Changes in mitochondrial morphology have been reported in the muscles of cachectic patients, however, so far, the underlying mechanisms are not clear. In our study, we have shown that mitochondrial morphology and lipid oxidation are closely linked. It is likely that an increased reliance on beta-oxidation of fatty acids in cachectic muscle causes it to exceed mitochondrial capacity. This in turn results in mitochondrial damage and dysfunction, and contributes to cancer-induced muscle wasting. Interestingly, in the muscles of mice and humans, it has been reported that fatty-acid beta-oxidation and mitochondrial dynamics can influence each other^46–48^, therefore, it would be interesting to explore the direct link between these two parameters in the future.

It has been reported that cachectic patients exhibit decreased dietary lipids and increased fatty acid release from the adipose tissue^49^. It is therefore likely that the depletion of available energy substrates (glycogen and fatty acids) contributes to the mitochondrial dysfunction and loss of muscle integrity. In this study, we have shown that coconut oil supplementation increases mitochondrial function and improves muscle integrity. While the underlying mechanism is so far unclear, we think this dietary supplementation most likely acts to increase lipid stores in general, therefore delaying mitochondrial damage caused by excessive beta-oxidation. However, it is also possible that a high-fat diet could first lead to reduced mitochondrial size^47,48^, which in turn can cause decreased beta-oxidation^46^, therefore, leading to a slower utilisation of lipids.

We reported a decrease in LD number in the muscles of both cachectic flies and mice. In contrast with this, studies in cachectic patient muscle samples have previously reported an accumulation of intramyocellular lipids^42,43^. When we examined LD levels in the flies, we did not measure intramyocellular lipid levels, as our images were taken longitudinally. Upon examination of LD levels in the muscles of the C-26 mouse model (via transverse section), we saw a decrease in extramyocellular LD levels, and an increase in intramyocellular LDs. It is therefore possible that the depletion of LDs we observed is in fact a mobilisation of extramyocellular/subsarcolemmal LDs to intramyocellular LDs, for their use in mitochondrial beta-oxidation. This would be consistent with reports that in starved mouse embryonic fibroblasts, fatty acids stored in LDs undergo transfer to mitochondria for beta-oxidation^50^. In the future, it would be interesting to explore whether the increase in intramyocellular LDs in human cachexia muscle samples also correlates with a decrease in extramyocellular LDs, and if this is associated with changes in fatty acid beta-oxidation.

Together, our study demonstrates that targeting FOXO, mitochondrial fusion/beta-oxidation and replenishing lipid stores in cachectic animals can prevent muscle wasting in cachexia. Therefore, this work opens up new avenues for finding therapeutic targets to prevent or attenuate the progression of cancer cachexia in patients.

## Acknowledgements

We are grateful to Kieran Harvey, Donna Denton, Helena Richardson, Alex Gould, and Tatsushi Igaki for generous sharing of fly stocks and antibodies. We would like to thank Bloomington Drosophila Stock Center, Vienna Drosophila Resource Center and Developmental Studies Hybridoma Bank for fly stocks and antibodies. We would like to also thank OZDros for *Drosophila* quarantine, Peter MacCallum Cancer Institute Microscopy Core for technical assistance. We are grateful to Kellie Veen and Khanh Nguyen for assistance with illustrations, Yuchen Bai for technical assistance with mouse immunostaining experiments, Michelle Meier for assistance with mitochondrial quantifications and Daniel Bakopolous for intellectual input in this project. We would also like to thank Edel Alvarez and Khanh Nguyen for critical reading of the manuscript. This work is funded by the NHMRC Ideas Grant APP1182847 and the Peter MacCallum Cancer Foundation.

## Author contribution

C.D, N.A, S.G, R.B, B.P, K.M, G.L,T.S, S.S.M, S.M.J, A.R.J and L.Y.C conducted the experiments; C.D and L.Y.C wrote the manuscript.

## Methods

### *Drosophila* stocks and husbandry

The following stocks were used from the Bloomington *Drosophila* stock centre:

*Mef2-GAL4* (BL27391),

*MHC-GAL4* (BL55133),

*UAS-Cat.A* (BL24621)

*UAS-Drp1* (BL51647),

*UAS-foxo^RNAi^* (BL27656),

*UAS-foxo^RNAi^* (BL32427), (of the two RNAi’s, *foxo^RNAi^* (27656) showed the stronger effect and was used for all other experiments

*UAS-luciferase* (BL64774),

*UAS-marf^RNAi^* (BL31157),

*UAS-marf^RNAi^* (BL67158) (of the two RNAi’s, *marf^RNAi^* (67158) showed the stronger effect and was used for all the experiments)

*UAS-mCherry^RNAi^* (BL35785),

*UAS-Opa1^RNAi^* (BL32358),

*UAS-pGFP-mCherryAtg8a* (BL37749),

*UAS-S6K^CA^* (BL6914),

*UAS-Sod1* (BL24750),

*UAS-whd^RNAi^* (BL33635),

*ey-FLP1;act>CD2>GAL4,UAS-GFP*^4^.

The following stocks were obtained from the Vienna Drosophila Resource Centre: *UAS-Impl2^RNAi^* (v30931).

The following stocks were also used:

*_w1118_*,

*ey-FLP1;UAS-dlg RNAi,UAS-RasV12 /CyO, Gal80;act>CD2>GAL4,UAS-GFP,*

*Ey-FLP1; QUAS-Ras^V^*^12^*, QUAS-scrib^RNAi^/ CyOQS; MHCGal4, act>CD2>QF, UAS-RFP/TMBQS*^4^,

*Ey-FLP1; QUAS-Ras^V^*^12^*, QUAS-scrib^RNAi^/ CyOQS; mef2Gal4, act>CD2>QF, UAS-RFP/TMBQS*^4^,

*UAS-Atg1^RNAi^* (Donna Denton),

*UAS-Atg1 6A* (Donna Denton),

*UAS-dFOXO* (Kieran Harvey),

*UAS-GFP* (Kieran Harvey),

*UAS-GPx1* (Tatsushi Igaki),

*UAS-lacZ^RNAi^* (Kieran Harvey),

*UAS-lsd2* (Alex Gould),

*UAS-Ras^V^*^12^ (Helena Richardson),

*mCherryAtg8a/CyOGFP*,

*mCherryAtg8a/CyOGFP;mCherry^RNAi^/TM6cSb*,

*mCherryAtg8a/CyOGFP;ImpL2^RNAi^/TM6cSb*.

Fly stocks were reared on standard *Drosophila* media, adults were allowed to lay for 24 hr at 25 °C and the progeny was then moved to 29 °C. Experiments were conducted on animals lacking tumours at wandering stage, and on tumour-bearing animals on a specific number of days after egg lay as indicated throughout the methods.

### Dietary supplementation and nutrient restriction

Fly food supplemented with nicotinamide at a concentration of 1 g of nicotinamide per 1 kg of normal food was created as follows. 100 g of regular food was melted in a beaker using a microwave until there were no solid lumps. 100 mg of nicotinamide^44^ (Sigma-Aldrich, #N0636-100G) was dissolved in 1 ml of sterile water at room temperature, added to the melted fly food, and mixed thoroughly. The combined mixture was divided into ten fly vials with approximately 10 ml of food per vial. 100 g of regular food without nicotinamide was also melted and poured into ten vials to be used as control food. Both nicotinamide and regular food was left to cool at 4 °C overnight. To control for density in feeding, crosses were laid on regular food. On day 1 AEL, before the embryos hatched, 20-30 embryos were taken and placed on a piece of cardboard and placed into the control or nicotinamide food. The larvae that hatch on the food were then fed on the specified diets until dissection.

High-fat diet fly food was created as follows. 70 g of regular food was melted in a beaker using a microwave until there were no solid lumps. 30 g of coconut oil^45^ (Community Co Virgin Coconut Oil 450 ml) was melted in a separate beaker using a microwave until there were no solid lumps. The melted coconut oil was then poured into the beaker with the 70 g regular food and mixed thoroughly. The mixture was then left to cool slightly for 5-10 mins, then was mixed thoroughly again to prevent the food from separating from the coconut oil. This was repeated if the mixture appeared to separate again. The combined mixture was divided into ten fly vials with approximately 10 ml of food per vial. 100 g of regular food without coconut oil was also melted and poured into ten vials to be used as control food. Both high-fat food and regular food was left to cool at 4 °C overnight. A small piece of tissue paper (approximately 1 x 3 cm) was placed into the food of control and high-fat vials to soak up excess coconut oil. To control for density in feeding, crosses were laid on regular food. On day 1 AEL, before the embryos hatched, 20-30 embryos were taken and placed on a piece of cardboard and placed into the control or high-fat food. The larvae that hatch on the food were then fed on the specified diets until dissection.

For starvation/nutrient restriction, the larvae were fed on normal food for 60 hr. Approximately 20 larvae were transferred onto either normal food or a diet consisting of 1% agar in PBS at this time point. The animals were then either fed or starved for 24 hr before dissection.

### *Drosophila* immunostaining

For FOXO, MHC staining as well as phalloidin staining (muscle integrity), larvae were heat-killed ^4,51^, muscle fillets (prepared as previously described^4,51^) were then fixed for 20 min in PBS containing 4% formaldehyde and washed three times for 10 mins each with PBS containing 0.3% Triton-X (PBST-0.3). For Atg8a and LipidTOX^TM^ experiments, animals were dissected in cold 1x PBS, fixed for 45 mins and washes were performed in 1x PBS. For ATP5A staining, DHE and TMRE experiments, animals were dissected in cold *Drosophila* Schneider’s Medium. ATP5A samples were fixed for 20 min followed by three 10 min washes in PBST-0.3. For DHE and TMRE there was no fixation or wash steps before staining. Tissues were then stained as per the manufacturer’s specifications. All samples were imaged on an Olympus FV3000 confocal microscope. Muscle integrity and TMRE samples were imaged using a 10x objective lens. FOXO, MHC, Atg8a, LipidTOX^TM^, and DHE were imaged with a 40x objective lens. ATP5A was imaged with a combination of 40x and 63x lenses. All confocal samples were mounted in glycerol except for LipidTOX^TM^ and TMRE, which were mounted in 1 x PBS and 25 nM TMRE in *Drosophila* Schneider’s Medium, respectively. Within a given experiment, all images were acquired using identical settings. Primary antibodies used: dFOXO (Abcam, 1:100, #ab195977), MHC (DHSB, 1:10, #3E8-3D3), ATP5A (Abcam, 1:500, #ab14748). Secondary donkey antibodies conjugated to Alexa 488 and Alexa 555 (Molecular Probes) were used at 1:200. DAPI 405 (Abcam, #ab228549) was used at 1:10000, Phalloidin 647 (Abcam, #ab176759) was used at 1:10000, HCS LipidTOX^TM^ Red Neutral Lipid Stain (Invitrogen, #H34476) was used at 1:1000. MitoTracker^TM^ Green FM (Invitrogen, #M7514) was used at a concentration of 250 nm^52^. TMRE was used as previously described (Invitrogen, #T669, 100 nm)^53^. DHE staining was performed as previously described (10 mins DHE stain)^54^.

### Glycogen staining

Muscle fillets were dissected in 1% BSA in PBS, fixed in 4% formaldehyde in PBS for 20 min, and washed twice in 1% BSA in PBS. Periodic acid stain (PAS) was used as previously described^55^. The samples were mounted in glycerol and imaged on an Olympus BX53 Brightfield microscope using a 4x objective lens.

### Experimental animals (mice)

All experiments were approved by the Animal Ethics Committee of The University of Melbourne and conducted in accordance with the Australian code of practice for the care and use of animals for scientific purposes as stipulated by the National Health and Medical Research Council (Australia). Male Balb/c mice were obtained from the Animal Resources Centre (Canning Vale, Western Australia). All mice were housed in the Biological Research Facility under a 12:12 hr light-dark cycle, with water and standard laboratory chow available *ad libitum*.

### Mouse model of cancer cachexia

The procedures used to thaw and count the Colon-26 (C-26) cells used to inject mice has been previously described^56^. Twelve-week-old male Balb/c mice were anesthetized with isoflurane (induction, 3-4% oxygen-isoflurane at 0.5 L.min^-1^; maintenance, 2-3% at 0.5 L.min^-1^), such that they were unresponsive to tactile stimuli. Mice were then given a subcutaneous (*s.c.*) injection of 5 × 10^5^ C-26 cells suspended in 100 µl of sterilized phosphate buffered saline (PBS; n = 4) or 100 µl of sterilized PBS only (control; n = 4) and recovered from anaesthesia on a heat pad. After 17-25 days, when end-point criteria was met, mice were anaesthetized deeply with sodium pentobarbitone (Nembutal; 60 mg/kg; Sigma-Aldrich, Castle Hill, NSW, Australia) via intraperitoneal (*i.p.*) injection and the tibialis anterior (TA) muscles were carefully excised, blotted on filter paper, trimmed of tendons and any adhering non-muscle tissue and weighed on an analytical balance. The LTA muscle was mounted in embedding medium, frozen in thawing isopentane and stored at -80 °C for subsequent analyses. Mice were killed by cardiac excision while still anesthetized deeply.

### Mouse immunostaining

Serial sections (8 µm) were cut transversely through the TA muscle using a refrigerated (-20 °C) cryostat (CTI Cryostat; IEC, Needham Heights, MA, USA). Frozen mouse TA tissue was thawed at room temperature for 10 mins, then fixed in 4% neutral buffered formalin for 10 mins and washed twice for 5 mins in 1x PBS. Samples were then stained as per the manufacturer’s specifications. Samples were mounted in 1x PBS and imaged on an Olympus FV3000 confocal microscope with a 40x objective lens. All images were acquired using identical settings. HCS LipidTOX^TM^ Red Neutral Lipid Stain (Invitrogen, #H34476) was used at 1:1000.

### Image analysis

All images were quantified using FIJI^57^. FOXO intensity was normalised to DAPI, FOXO and DAPI levels were quantified by drawing a circle around the nucleus in the DAPI channel, and the mean grey value (m.g.v.) determined for FOXO and DAPI channels. To measure fluorescence intensity of MHC, Atg8a, PAS and DHE, a ROI was drawn around a sarcomere (MHC), nucleus plus adjacent to nucleus (Atg8a), section of a muscle segment (PAS), and nucleus only (DHE), on the z-plane where fluorescence was most intense. For the MHC, single Atg8a, PAS, and DHE quantifications, the levels of fluorescence were calculated with respect to background fluorescence, using total corrected cell fluorescence (TCCF), as described previously^58^. For the Tandem Atg8a-mCherry-GFP quantifications, a measurement was taken in both the mCherry and GFP channels, and total autophagy was calculated as a ratio of Atg8a-mCherry to Atg8a-GFP.

Percentage muscle/cuticle was determined using FIJI as previously described^4,51^. In brief, dissected muscle fillets stained with Phalloidin to mark actin were analysed using a FIJI macro^51^. A ROI was drawn around the cuticle of the muscle fillet, and the image was converted to a binary mask using the “Auto Threshold” tool. The total area of fluorescence detected within the ROI was divided by the total ROI area, which we calculated as % muscle attachment.

The number of LDs present in the fly muscle was determined through the use of a macro in FIJI (Supplementary File 1). In brief, files were imported into FIJI, and a 200 x 200 pixel ROI was created on an 8-bit converted representative slice. The image was cropped to the ROI, then the “Auto-threshold” function was used to convert the image into binary. The “Analyse Particles” function using a size range of “0.00-10” was then used to count the number of LDs present in the image.

The number of large extramyocellular LDs present in the mouse muscle was determined using a macro in FIJI (Supplementary File 1). In brief, files were imported into FIJI, a representative slice was converted to 8-bit. The “Auto-threshold” function was used to convert the image into binary. The “Analyse Particles” function using a size range of “2-infinity” was then used to count the number and size of extramyocellular LDs present in the image.

The number of intramyocellular LDs present in the mouse muscle was determined using a macro in FIJI (Supplementary File 1). In brief, files were imported into FIJI, and the “Auto-threshold” function was used to convert the image into binary. Five polygon ROIs that each encompassed the interior of a different myofiber was created on an 8-bit converted representative slice. The “Analyse Particles” function using a size range of “0.00-10” was then used to count the number and size of LDs present in the image. The density and size of LDs was averaged between the five myofibers for one image.

The proportion of large mitochondria in the muscle was determined through the use of a macro in FIJI (Supplementary File 2). For more details on the use of the macro and how large mitochondria were selected, see Figure S1. The output of the macro was a list of individual mitochondrial areas. Once the output was acquired, a Log_10_ transformation was then applied to this list of areas to bring it closer to a normal distribution. The data was then binned into four sizes, those with a Log_10_ transformed value: X ≤ -0.5, -0.5 < X ≤ 0, 0 < X ≤ 0.5, and 0.5 < X ≤ 1.0. The mitochondria in the bin of 0.5 < X ≤ 1.0 were considered the largest mitochondria, and the proportion of these out of the total number of mitochondria was calculated.

The level of TMRE activity was determined through the use of a macro in FIJI (Supplementary File 3). In brief, files were imported into FIJI and split into MitoTracker^TM^ Green and TMRE channels. A max intensity z-projection of the MitoTracker^TM^ Green channel was converted to 8-bit, smoothed using the “Gaussian Blur” and “Remove Outliers” functions, and a square ROI was drawn around the muscle fillet. The image was then converted to a binary mask using the “Auto Threshold” function using the “IsoData” method. A selection was then created around all the thresholded MitoTracker^TM^ Green area, and this area was measured. A max intensity z-projection of the TMRE channel was then processed in the same way as the MitoTracker^TM^ Green channel. The final result was an area measurement for total mitochondria via MitoTracker^TM^ Green and active mitochondria via TMRE. The total area of TMRE was divided by the total area of MitoTracker^TM^ Green, to give a percentage value of how many total mitochondria are active.

### Electron Microscopy

*Drosophila* 3^rd^ instar larvae were dissected and fixed in 2.5% glutaraldehyde solution in 0.1 M sodium cacodylate buffer overnight at 4 °C or for 2 hr at room temperature. The samples were then washed with 0.1 M sodium cacodylate, followed by staining with 1% osmium tetroxide and 2% uranyl acetate using a Pelco Biowave and washed in 0.1 M sodium cacodylate. They were then dehydrated in an ethanol series (1x-50%, 70%, 90% and 2x-100%) followed by infiltration with increasing concentrations of epon resin (1x-25%, 50%, 75% and 2x–100%). Samples were subsequently processed in the resin with the Biowave high vacuum function before being embedded in fresh resin and polymerized in a 60 °C oven for 48 hr. Formvar-coated, one-slot grids were used to collect thin sections (50 nm) obtained via a Leica Ultracut UC6 Ultramicrotome by taking longitudinal sections of muscles. Images were collected in a JEOL 1011 electron microscope at 80 kV.

### Proteomics

Samples were lysed by tip-probe sonication in 1% SDS containing 10 mM tris(2-carboxyethyl)phosphine and 40 mM chloroacetamide in 100 mM HEPES pH 8.5. The lysate was incubated at 95 °C for 5 mins and centrifuged at 20,000 x g for 30 mins at 4 °C. Peptides were prepared using a modified SP3 approach with paramagnetic beads^59^. Briefly, lysates were shaken with a 1:1 mixture of hydrophilic and hydrophobic Sera-Mag SpeedBeads (GE Healthcare) in a final concentration of 50% ethanol for 8 mins at 23 ℃. The beads were washed three times with 80% ethanol and dried at 23 ℃ for 20 mins. Proteins were digested directly on the beads in 50 µL of 10% trifluoroethanol in 100 mM HEPES, pH 7.5 sequencing-grade LysC (Wako Chemicals) and sequencing-grade trypsin (Sigma) for 16 hr at 37 ℃. The supernatant containing peptides were removed and mixed with 150 µl of 1% trifluoroacetic acid (TFA) and purified using styrenedivinylbenzene-reverse phase sulfonate microcolumns. The columns were washed with 100 µl of 99% isopropanol containing 1% TFA followed by 100 µl of 99% ethyl acetate containing 1% TFA followed by 5% acetonitrile containing 0.2% TFA and eluted with 80% acetonitrile containing 1% ammonium hydroxide then dried by vacuum centrifugation. Peptides were resuspended 2% acetonitrile, 0.1% TFA and stored at - 80 ℃.

Peptides were separated on a 40 cm x 75 µm inner diameter PepMap column packed with 1.9 µm C18 particles (Thermo Fisher) using Dionex nanoUHPLC. Peptides were separated using a linear gradient of 5 – 30% Buffer B over 70 mins at 300 nl/min (Buffer A = 0.1% formic acid; Buffer B = 80% acetonitrile, 0.1% formic acid). The column was maintained at 50 °C coupled directly to an Orbitrap Exploris 480 mass spectrometer (MS). A full-scan MS1 was measured at 60,000 resolution at 200 m/z (350 – 951 m/z; 50 ms injection time; 2.5e6 automatic gain control target) followed by data-independent analysis (16 m/z isolation with 37 windows and a 1 m/z overlap, 28 normalized collision energy; 30 K resolution; auto injection time, 2e6 automatic gain control target).

Mass spectrometry data were processed using Spectronaut DirectDIA (v15.1.210713.50606) and searched against the Drosophila melanogaster UniProt database (October, 2019) using all default settings with peptide spectral matches and protein false discovery rate (FDR) set to 1%. The data were searched with a maximum of 2 miss-cleavages, and methionine oxidation and protein N-terminus acetylation were set as variable modifications while carbamidomethylation of cysteine was set as a fixed modification. Quantification was performed using MS2-based extracted ion chromatograms employing 3-6 fragment ions >450 m/z with automated fragment-ion interference removal as described previously^60^. Data was analysed in Perseus^61^ and included median normalisation and differential expression analysis using t-tests with multiple hypothesis correction using Benjamini-Hochberg FDR adjustment.

### Human sample collection

Collection of biospecimens of rectus abdominus muscle from patients with pathologically diagnosed pancreatic ductal adenocarcinoma (PDAC; n = 10 females/10 males) undergoing tumour resection surgery was compliant with an approved Institutional Review Board protocol at the University of Florida, with written informed consent obtained from all patients, and conformed to the Declaration of Helsinki. The detailed patient demographics and analysis of preoperative CT scans for skeletal muscle index (SMI) and muscle radiation attenuation as quantitative measures of skeletal muscularity and myosteatosis, respectively, have been published previously^41,62^. The methods for obtaining the microarray data have been published previously^41^.

### Statistical analysis

All statistical analyses were conducted using GraphPad Prism 9.0 (©GraphPad software Inc.). For experiments measuring % muscle/cuticle, FOXO/DAPI levels, PAS staining, MHC staining, DHE staining, Atg8a intensity, and TMRE/MitoTracker^TM^ Green ratios, experimental values were normalised to the average value of their respective controls. At least three animals per genotype were used for all muscle experiments. For FOXO, DHE, Atg8a, MHC staining intensity quantifications, individual data points represent fluorescence intensity of a single nucleus or muscle fibre. For muscle integrity, PAS staining, LD number, mitochondrial size, and TMRE quantifications, individual data points represent a single larva. For experiments with two genotypes or treatments, two-tailed unpaired student’s t-tests were used to test for significant differences. The Welch’s correction was applied in cases of unequal variances, and the Mann-Whitney U tests was used in the cases of violated normality. For experiments with more than two genotypes, significant differences between specific genotypes were tested using a one-way ANOVA and a subsequent Šidák post-hoc test. A Brown-Forsythe correction was applied in cases of unequal variances, and in the cases of violated normality, the Kruskal-Wallis test was used. The results for all post-hoc tests conducted in a given analysis are shown in graphs. For all graphs, error bars represent SEM. p and adjusted-p values are reported as follows: p>0.05, ns (not significant); p<0.05, *; p<0.01, **; p<0.001, ***; p<0.0001, ****.

---

**Figure S1.**
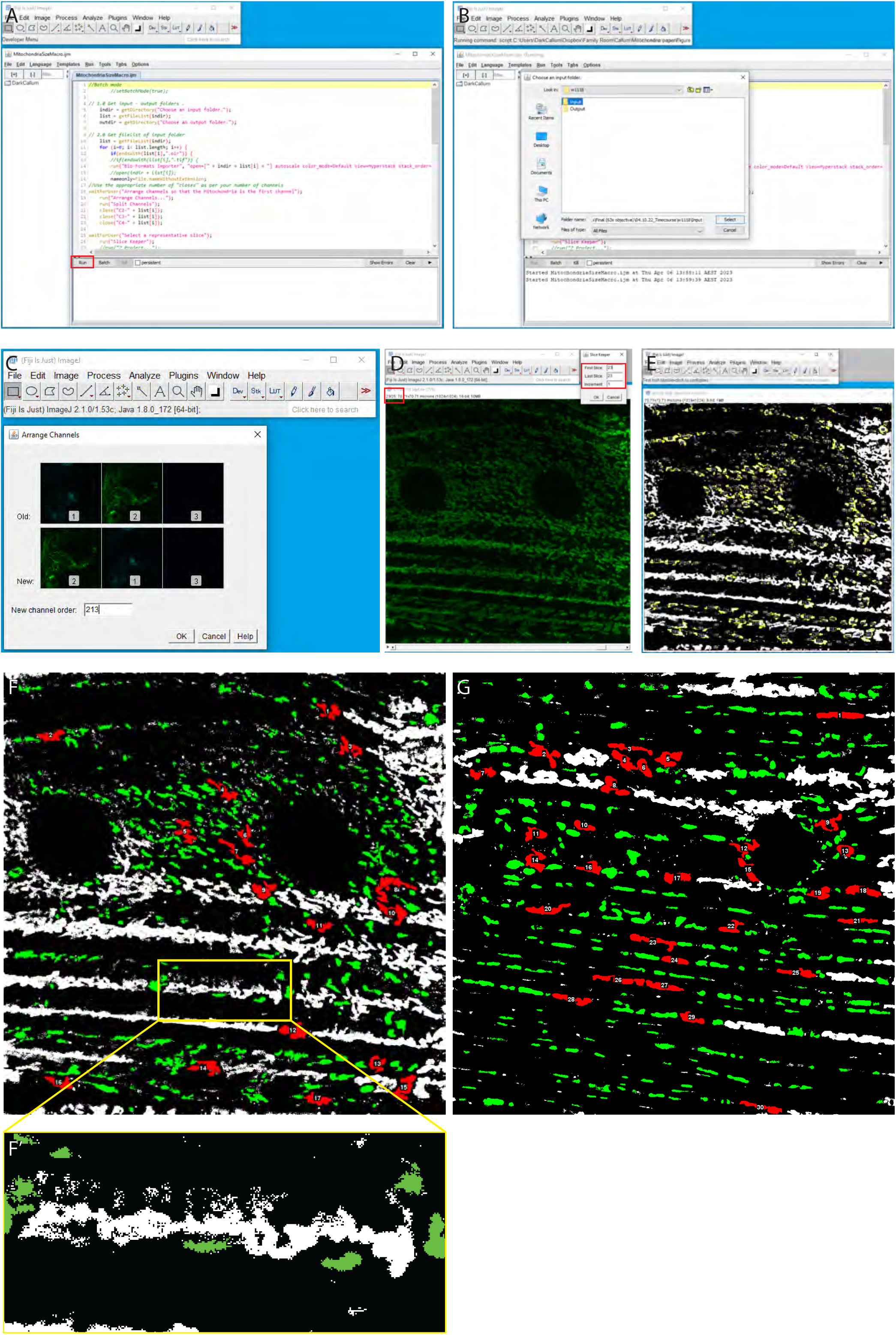
Instructions for utilising FIJI macro for the quantification of mitochondrial size. A) Import the Mitochondrial Macro (Supplementary File 2) into FIJI and hit “Run”, as indicated by the red box. B) The macro will ask you to select first an Input folder and then an Output folder. We recommend creating separate folders and then moving the z-stacks you have acquired and require to be analysed into the “Input” folder. C) Next, the macro will seek to isolate the fluorescence channel for the mitochondrial marker (in this case ATP5A). To do this, it will ask "Arrange channels so that the Mitochondria is the first channel". Click “Ok”. When the “Arrange Channels” window appears you will see the original channel order, in this case the mitochondria channel was the second channel out of three (Order: 123), so we have changed the order to put it first, by typing in the “New channel order” box (Order: 213). The macro then splits the channels and closes all other channels. D) You then will be asked to “Select a representative slice”. Before you click “Ok”, scroll through the z-stack to find a slice that best shows the sub-sarcolemmal mitochondria. Here, we have selected slice 23, as indicated by the red boxes. Click “Ok”, then the “Slice Keeper” window will appear. Type your desired slice number into both the “First Slice” and “Last Slice” boxes and put the number “1” into the “Increment” box. It will then perform an 8-bit conversion, change the colour to Grays, and will “Auto Threshold” the image using the “IsoData” method. E) The macro will then use the “Analyze Particles” tool to measure the size and number of mitochondria in the image. We have set the macro to only measure structures been 0.2 and 7 μm^2^ in size. This avoids detecting structures that are too small (< 0.2 μm^2^), which are likely background, and avoids detecting structures that are too big (> 7 μm^2^). The macro will then output the thresholded image, as well as an excel file with the number of mitochondria that were measured and their respective sizes. It will then move onto the next z-stack. F, F’) An example of mitochondria that are above the “large” threshold (log_10_ transformed value, 0.5 < X ≤ 1.0, depicted in red), and below the threshold (log_10_ transformed value, X ≤ 0.5, depicted in green), in a control muscle fillet. All other mitochondria depicted in white were not measured as they were outside the “Analyze Particles” size range. 17 mitochondria were considered in the “large” range (for more details on how mitochondria were binned for size, see Methods and Materials. The yellow box shows a zoomed in example of a large structure (marked in white) that is excluded from the analysis in F’, as it exceeds the size threshold. G) An example of mitochondria that are above the “large” threshold (log_10_ transformed value, 0.5 < X ≤ 1.0, depicted in red), and below the threshold (log_10_ transformed value, X ≤ 0.5, depicted in green), in a 7 days AEL *Ras^V12^dlg^RNAi^* muscle fillet. All other mitochondria depicted in white were not measured as they were outside the “Analyze Particles” size range. 30 mitochondria were considered in the “large” range.

**Figure S2.**
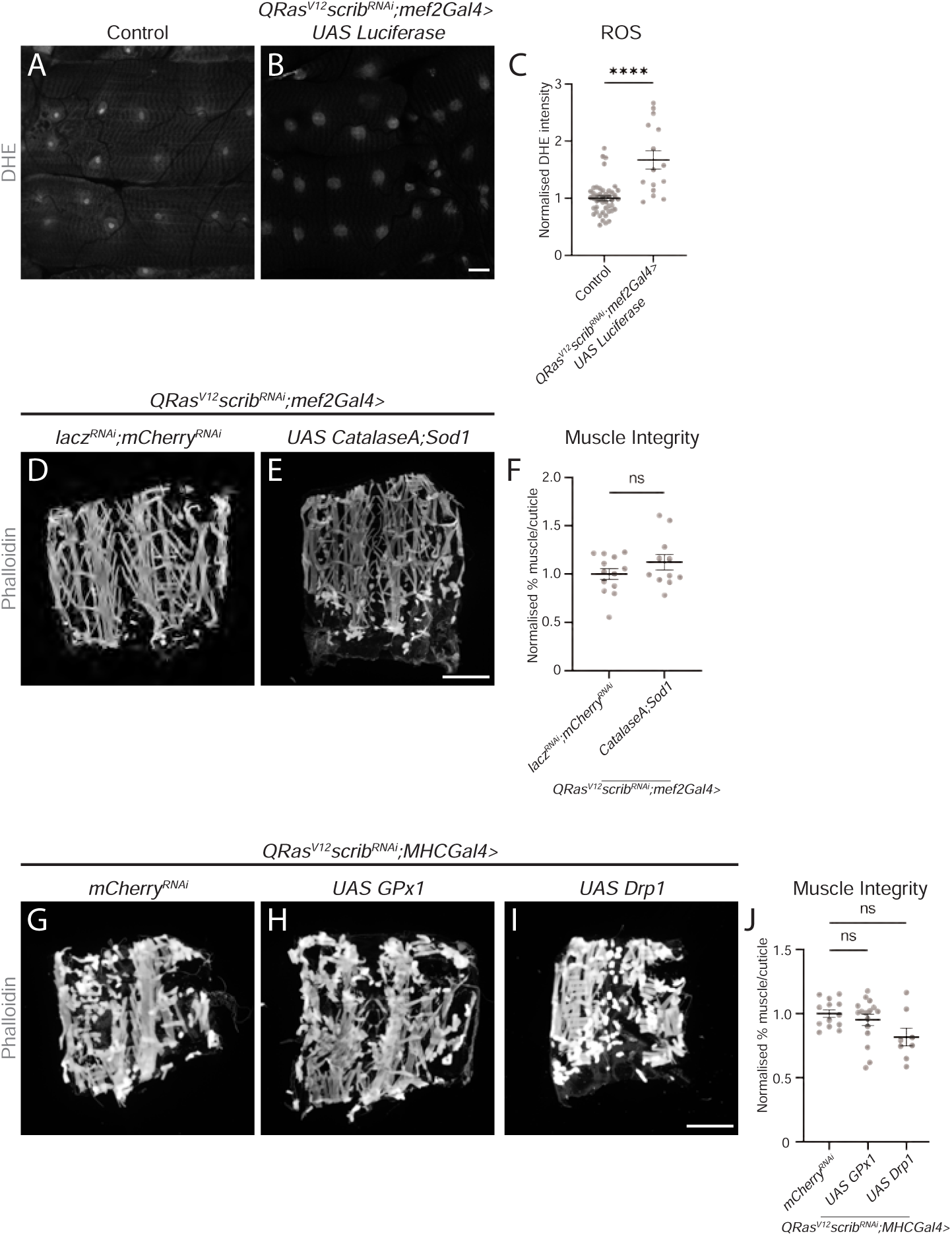
Reduced ROS in the muscles of tumour-bearing animals does not rescue muscle wasting. A, B) Representative images of DHE staining in the muscles of control and *QRas^V12^scrib^RNAi^;mef2-Gal4>UAS luciferase* larvae. C) Quantification of DHE staining in A and B, performed using Mann-Whitney U (n = 45, 15). D, E) Representative muscle fillets from *QRas^V12^scrib^RNAi^;mef2-Gal4>lacz^RNAi^;mCherry^RNAi^*and *QRas^V12^scrib^RNAi^;mef2-Gal4>UAS-CatalaseA;UAS-Sod1* larvae, stained with Phalloidin to visualise actin. F) Quantification of muscle integrity in D and E, performed using Student’s t-test (n = 13, 11). G, H, I) Representative muscle fillets from *QRas^V12^scrib^RNAi^;MHC-Gal4>mCherry^RNAi^* (G), *QRas^V12^scrib^RNAi^;MHC-Gal4>UAS-GPx1* (H) and *QRas^V12^scrib^RNAi^;MHC-Gal4>UAS-Drp1* (I) larvae, stained with Phalloidin to visualise actin. J) Quantification of muscle integrity in G, H and I, performed using Kruskal-Wallis as part of an analysis with S3 R, which used the same controls (n = 13, 16, 8). Images were taken at 6 days AEL for (A), and 7 days AEL for (B, D, E, G, H and I). Scale bars: 20 μm for (A, B), and 500 μm for (D, E, G, H and I). All error bars are +/- SEM. P values are: ns (not significant), p > 0.05; *, p < 0.05; **, p < 0.01; ***, p < 0.001; ****, p < 0.0001.

**Figure S3.**
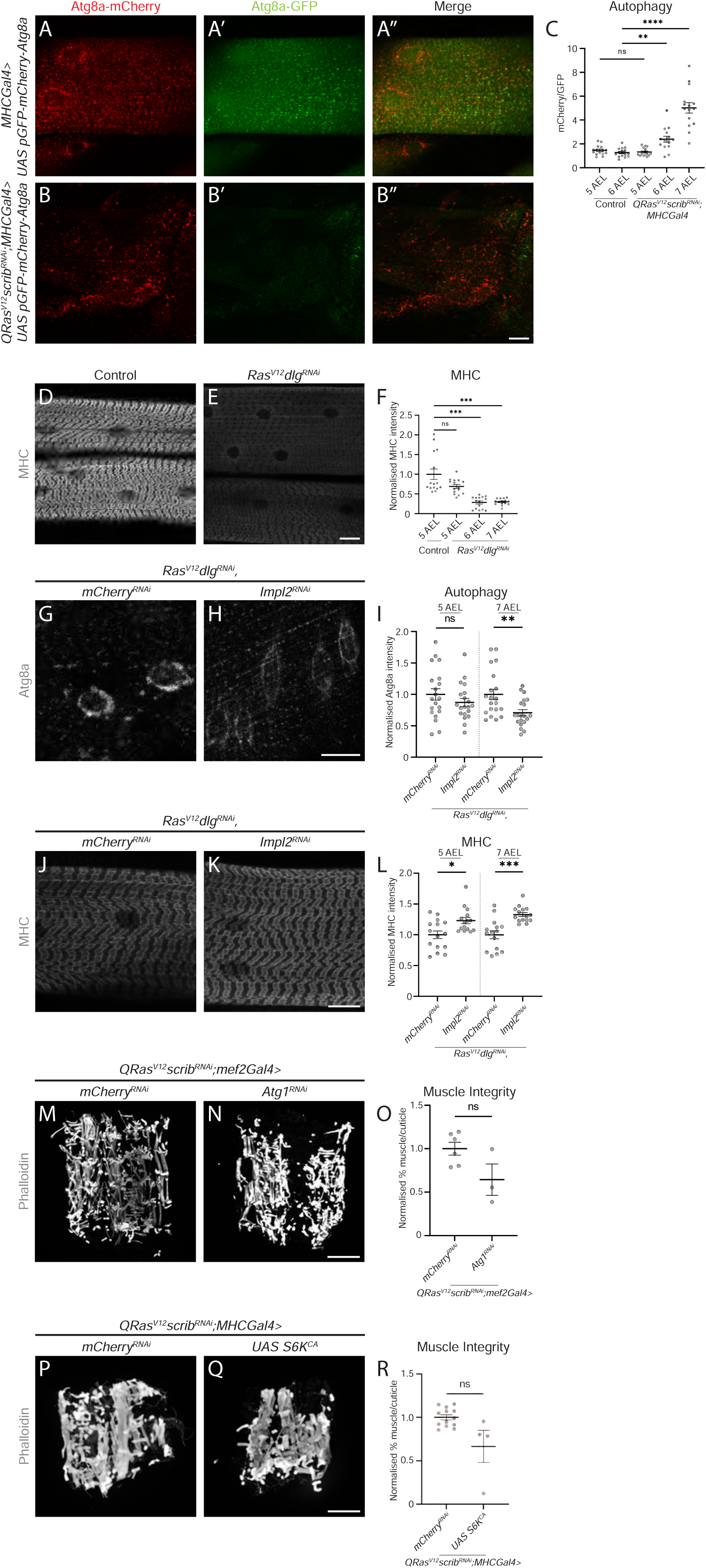
Muscles of tumour bearing *Drosophila* larvae show defects associated with autophagy and translation, but these phenotypes are not necessary for muscle wasting. A, A’, A’’, B, B’, B’’) Representative images of larval muscles of *MHCGAl4* and *QRas^V12^scrib^RNAi^;MHC-Gal4* animals crossed to a reporter of autophagy, Atg8a, tagged with both mCherry (A, B) and GFP (A’, B’). Merged images are shown in A’’ and B’’. C) Quantification of the ratio of Atg8a-mCherry to Atg8a-GFP in control and *Ras^V12^dlg^RNAi^* larvae at days 5-7 AEL, performed using Brown-Forsythe (n = 15, 15, 15, 15, 15). D, E) Representative images of Myosin Heavy chain (MHC) staining in the muscles of control and *Ras^V12^dlg^RNAi^* larvae. F) Quantification of MHC staining in control and *Ras^V12^dlg^RNAi^* larvae from days 5-7 AEL, performed using Brown-Forsythe (n = 15, 15, 15, 15). G, H) Representative images of larval muscles of *Ras^V12^dlg^RNAi^,mCherry^RNAi^*and *Ras^V12^dlg^RNAi^,ImpL2^RNAi^* crossed to a reporter of autophagy, Atg8a, tagged with mCherry. I) Quantification of Atg8a-mCherry levels in G and H, as well as from earlier timepoints, performed using Student’s t-test (5 days AEL), Welch’s t-test (6 days AEL), and Mann-Whitney U (7 days AEL, n = 20, 20, 20, 20, 20, 20). J, K) Representative images of Myosin Heavy chain (MHC) staining in the muscles of *Ras^V12^dlg^RNAi^,mCherry^RNAi^* and *Ras^V12^dlg^RNAi^,ImpL2^RNAi^* larvae. L) Quantification of MHC staining in J and K, as well as staining from earlier timepoints, performed using Mann-Whitney U (5 and 6 days AEL) and Welch’s t-test (7 days AEL, n = 15, 15, 15, 15, 15, 15). M, N) Representative muscle fillets from *QRas^V12^scrib^RNAi^;mef2-Gal4>mCherry^RNAi^*and *QRas^V12^scrib^RNAi^;mef2-Gal4>Atg1^RNAi^* larvae, stained with Phalloidin to visualise actin. O) Quantification of muscle integrity in M and N, performed using Student’s t-test (n = 6, 3). P, Q) Representative muscle fillets from *QRas^V12^scrib^RNAi^;MHC-Gal4>mCherry^RNAi^*, *QRas^V12^scrib^RNAi^;MHC-Gal4>UAS-S6K^CA^* larvae, stained with Phalloidin to visualise actin. R) Quantification of muscle integrity in P and Q, performed using Kruskal-Wallis as part of an analysis with S2 J, which used the same controls (n = 13, 4). Images were taken at 5 days AEL for (A, A’, A’’ and D), and 7 days AEL for (B, B’, B’’, E, G, H, J, K, M, N, P and Q). Scale bars: 20 μm for (A, A’, A’’, B, B’, B’’, D, E, G, H, J and K), and 500 μm for (M, N, P and Q). All error bars are +/- SEM. P values are: ns (not significant), p > 0.05; *, p < 0.05; **, p < 0.01; ***, p < 0.001; ****, p < 0.0001.

**Figure S4.**
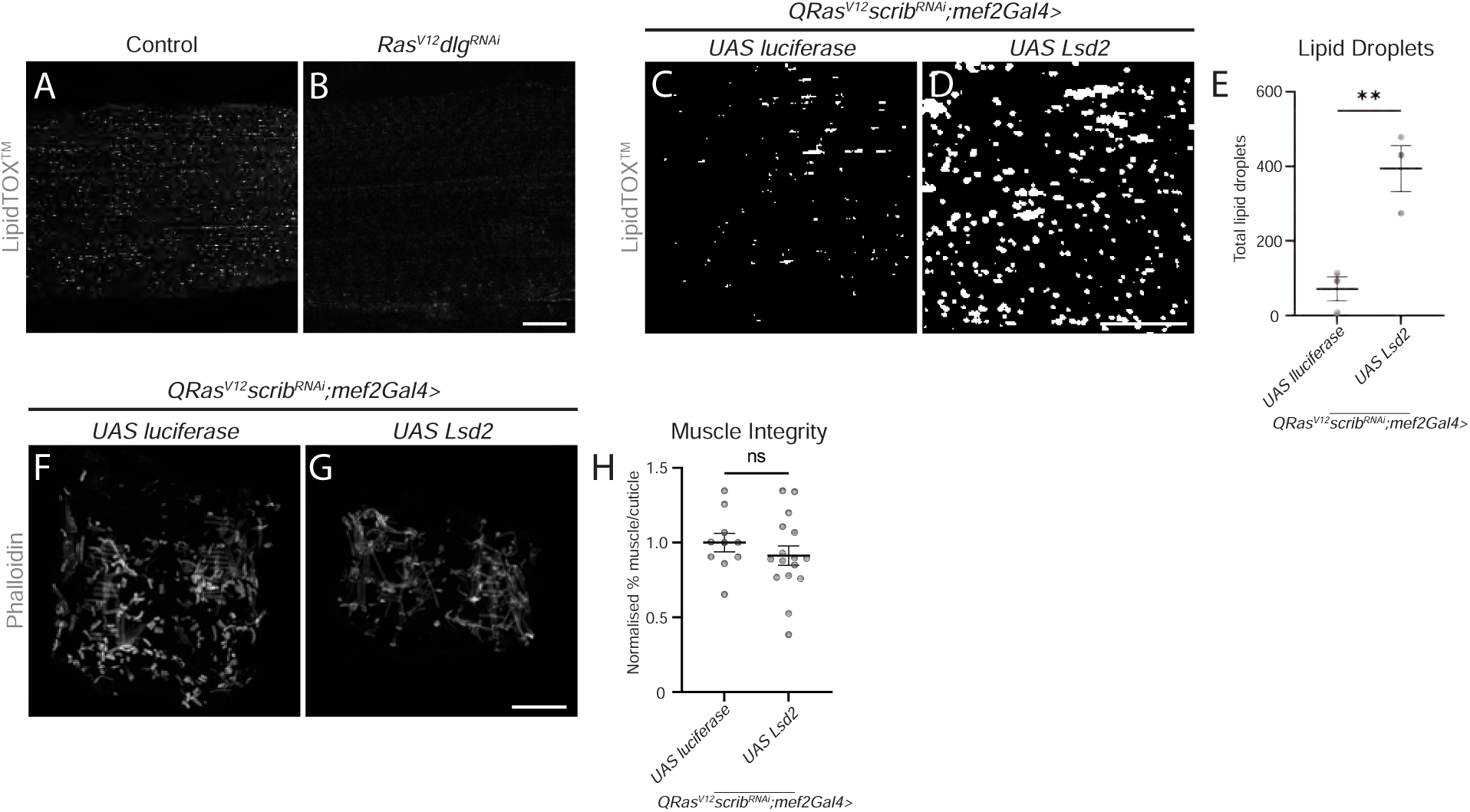
Increasing storage of lipids in lipid droplets does not rescue muscle integrity. A, B) Representative images of lipid droplets (LDs) stained with LipidTOX^TM^ in the muscles of control and *Ras^V12^dlg^RNAi^*larvae. C, D) Binary representation of LDs stained with LipidTOX^TM^ in the muscles of *QRas^V12^scrib^RNAi^;mef2-Gal4>UAS-luciferase* and *QRas^V12^scrib^RNAi^;mef2-Gal4>UAS-lsd2* larvae. E) Quantification of the number of LDs in C and D, performed using Student’s t-test (n = 3, 3). F, G) Representative muscle fillets from *QRas^V12^scrib^RNAi^;mef2-Gal4>UAS-luciferase* and *QRas^V12^scrib^RNAi^;mef2-Gal4>UAS-lsd2* larvae, stained with Phalloidin to visualise actin. H) Quantification of muscle integrity in N and O, performed using Student’s t-test (n = 10, 16). Images were taken at 5 days AEL for (A), 6 days AEL for (C and D), and 7 days AEL for (B, F and G). Scale bars: 20 μm for (A, B, C and D), and 500 μm for (F and G). All error bars are +/- SEM. P values are: ns (not significant), p > 0.05; *, p < 0.05; **, p < 0.01; ***, p < 0.001; ****, p < 0.0001.

